# Protein codes promote selective subcellular compartmentalization

**DOI:** 10.1101/2024.04.15.589616

**Authors:** Henry R. Kilgore, Itamar Chinn, Peter G. Mikhael, Ilan Mitnikov, Catherine Van Dongen, Guy Zylberberg, Lena Afeyan, Salman Banani, Susana Wilson-Hawken, Tong Ihn Lee, Regina Barzilay, Richard A. Young

## Abstract

Cells have evolved mechanisms to distribute ∼10 billion protein molecules to subcellular compartments where diverse proteins involved in shared functions must efficiently assemble. Here, we demonstrate that proteins with shared functions share amino acid sequence codes that guide them to compartment destinations. A protein language model, ProtGPS, was developed that predicts with high performance the compartment localization of human proteins excluded from the training set. ProtGPS successfully guided generation of novel protein sequences that selectively assemble in targeted subcellular compartments. ProtGPS also identified pathological mutations that change this code and lead to altered subcellular localization of proteins. Our results indicate that protein sequences contain not only a folding code, but also a previously unrecognized code governing their distribution in specific cellular compartments.

## Introduction

Groups of proteins involved in shared functions must efficiently assemble to fulfill their physiological functions^1^. For example, the fidelity of gene transcription hinges on the assembly of over a hundred different proteins at promoters, where some bind DNA sequences directly and others interact with DNA-bound proteins instead^2,3^. Selective protein-protein and protein-nucleic acid interactions are thought to be the predominant driving force leading to the assembly of specific proteins at locations where they carry out diverse functions^4-7^. Shape complementarity among structurally stable portions of proteins have dominated models of protein assembly, but there is now considerable evidence that large assemblies of proteins with shared functions also occur through weak multivalent noncovalent interactions^8-15^. Nearly all cellular functions involve formation of such assemblies, which have been described as condensates, aggregates, puncta, hubs and non-membrane bound compartments (Fig. 1A). In a recent study, we used small chemical probes to demonstrate that different condensates can harbor distinct internal chemical environments, suggesting that such assemblies have different solvent properties^16^. It is thus possible that protein molecules that assemble selectively with others in a condensate do so, in part, as a consequence of their compatibility with the internal solvating environment of that compartment^17-20^. Integration of contributions from specific interactions (e.g., DNA-protein binding, protein-protein interactions) and nonspecific interactions (e.g., transient noncovalent interactions) is challenging to model, but protein language models provide a mechanism for generally incorporating diverse contributions. If such a protein language model could be developed, it would have important implications for our understanding of cellular function and dysfunction by providing evidence of a protein code embedded in amino acid sequences guiding distribution.

**Fig 1.**
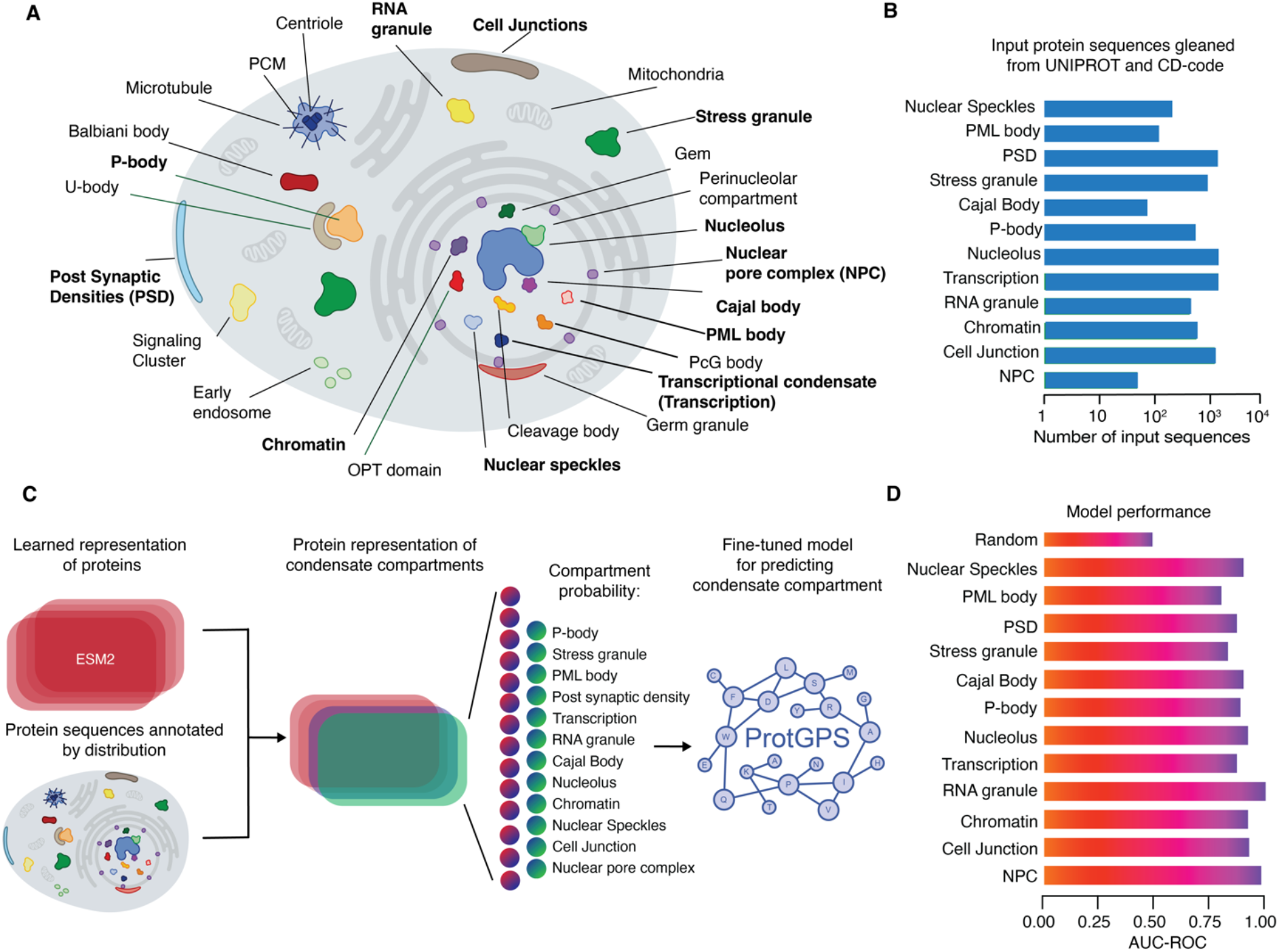
ProtGPS classifies protein compartment with high performance. **A**. Graphical depiction of some cellular compartments found in eukaryotic cells, compartments in bold were studied in this work. **B**. Bar graph showing the number of protein sequences gathered from UNIPROT and the CD-code database used in the development of ProtGPS. **C**. Schematic showing the approach toward developing ProtGPS. **D**. Bar graph showing the area under the receiver-operator curve for classification of withheld test data (15 % of total) with ProtGPS.

### Evidence for shared protein codes in condensate compartments

To learn whether collections of proteins that assemble into specific condensate compartments have shared protein codes, we adapted an evolutionary scale protein transformer language model (ESM2) to predict protein assembly into distinct compartments^21,22^. The transformer architecture of ESM2 allows for simultaneous relationships between all amino acids in an input sequence to be learned, providing a general strategy to detect protein codes embedded in the amino acid sequence of a protein. We focused our studies on a set of 5,541 human protein sequences that have been annotated for twelve condensate compartments using the UNIPROT^23^ and CD-Code^24^ databases (Fig. 1B). The compartment identities of the proteins in these databases were determined with various experimental techniques and curated by experts in compartment annotation and whole sequences were used as input^44^. Compartment annotated whole protein sequences were used as input. A neural network classifier was jointly trained with ESM2 to develop a model, termed ProtGPS, which computes the independent probability of a protein being found within each of the twelve different condensate compartments (Fig. 1C). The area under the receiver operator curve (AUC-ROC) showed that protein compartments could be predicted with remarkable accuracy (0.83-0.95) across the 12 different compartments (Fig. 1D). The performance of the ProtGPS model indicates it detects patterns in the protein primary structure that differentiates these condensate compartments.

### Guided generation of novel protein sequences for compartment selectivity

In order to validate that ProtGPS has learned the protein codes associated with condensate localization, we sought to design novel protein sequences that, when produced in cells, would selectively assemble into a compartment of interest. To test this idea, we initially designed protein sequences using an autoregressive greedy search algorithm (GS)^25^. We repeatedly feed a growing sequence into ProtGPS and extend the sequence with additional residues at the N-terminus up to a desired length, choosing an amino acid at each step that causes a protein to be classified as a match for the desired compartment (Fig. S1). For each protein, a plasmid was constructed that encoded a generated polypeptide of up to 150 amino acids with an N-terminal nuclear localization sequence to ensure transport into the nucleus and a C-terminal mCherry protein to ascertain the location of the protein by microscopy. In all, we generated eight novel protein sequences were designed to assemble selectively into nucleoli (Table S1). However, although these proteins entered the nucleus, they failed to assemble selectively into nucleoli (Fig. S1).

The failure of our initial efforts to generate proteins that selectively compartmentalize in nucleoli led us to consider how the GS algorithm might be imperfect for this task and motivated the design of another approach that might be more successful. With GS and ProtGPS, protein sequences are generated without consideration of the chemical space of proteins found in nature, but are over optimized towards prediction of subcellular compartments. We sought to create an approach that could overcome this limitation by applying a concept borrowed from medicinal chemistry, where it is common to consider whether a molecule shares desirable physicochemical properties with others^26,27^. We also chose to favor the use of “intrinsically disordered” domains because they have been implicated in protein association with condensates and because they are less likely to introduce competing folded states^28,29^. To apply these concepts toward protein generation, we sought to constrain generation to (1) proteins in the chemical space^30^ learned by ESM2, (2) domains that are intrinsically disordered, and (3) sequences that should localize to the intended compartment. Thus, we used additional features of protein chemical space and intrinsic disorder for our Markov Chain Monte Carlo algorithm (MCMC) (Fig. 2A).

**Fig. 2.**
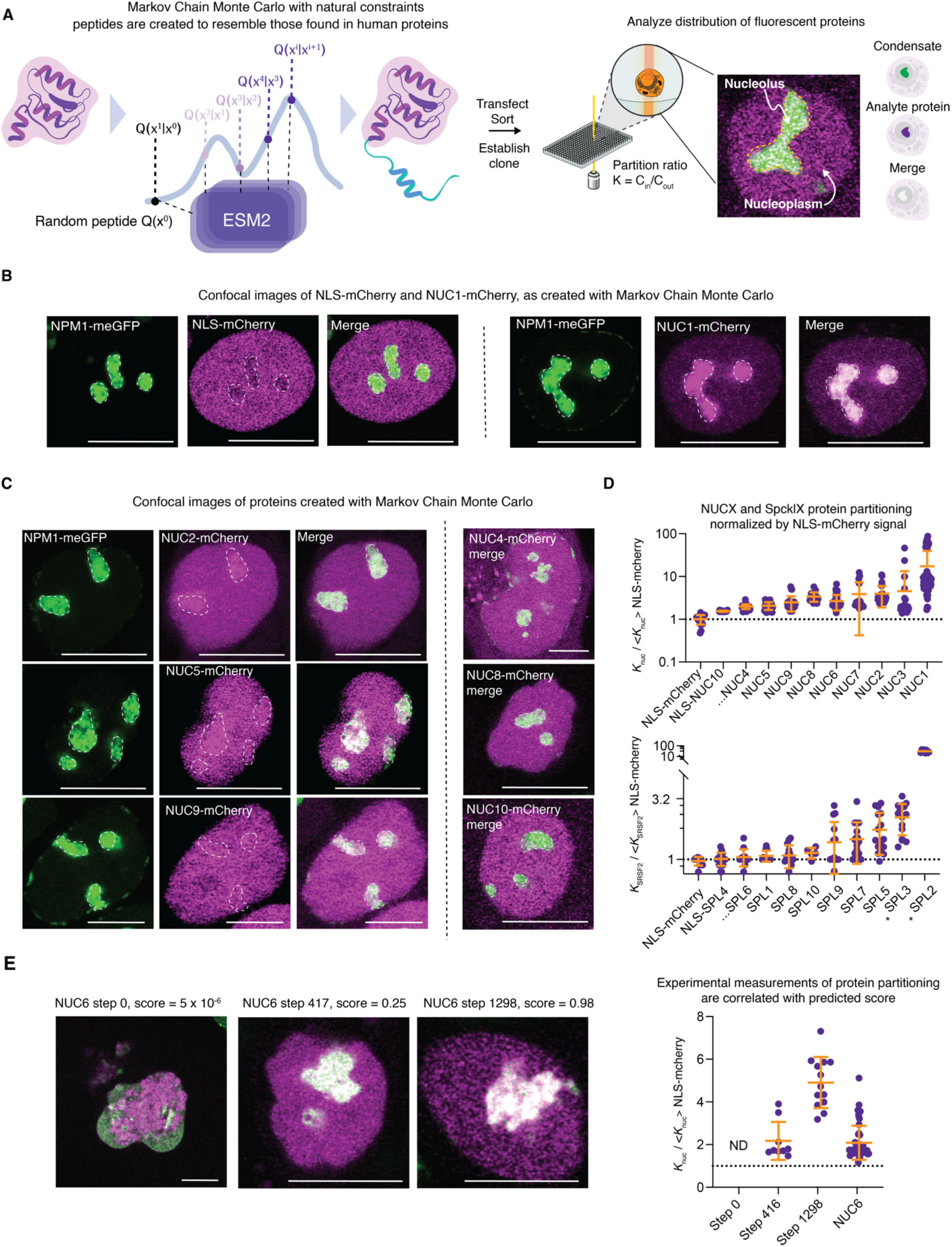
Generative modeling creates novel proteins that concentrate in a desired condensate. **A**. Schematic showing the use of Naturally constrained Markov Chain Monte Carlo to generate proteins and assay them in live cells (MCMC) (*see supporting information for more details*). **B**. Live cell image of a colon cancer cell (HCT-116) tagged at the endogenous NPM1 locus with GFP and expressing nucleolus targeted protein NUC1-mCherry, scale: 10 microns. **C**. Live cell confocal micrographs of NUCX-mCherry proteins in HCT-116 cells expressing NPM1-GFP from the endogenous locus cells, scale: 10 microns. **D**. Dot plots showing the measured partition ratios of NUCX (*K*_*x*_ =I_nucleolus_ / I_nucleoplasm_) and SPLX-mCherry (*Kx* = I_SRSF2_ / I_nucleoplasm_ or = I_SRSF2_ / I_cytoplasm_, as indicated by *) proteins relative to the NLS-mCherry control protein, dotted line is the average value of NLS-mCherry protein. See Table S3 for more information. **E**. Live cell images and quantification showing the relationship of measured partition ratios (*K*_*x*_ = I_nucleolus_ / I_nucleoplasm_) into the nucleolus by proteins on the NUC6-mCherry trajectory to its computed probability of partitioning.

We then used MCMC to perform guided generation of proteins, using the additional features described above, that would selectively assemble into two condensate assemblies, nucleoli^9^ and nuclear speckles^31^, that were selected because they are well-studied, have distinctive functions and morphologies, and possess unambiguous marker proteins (Fig. 2A). A total of twenty 100 amino acid long protein sequences, ten targeted to nucleoli and ten to nuclear speckles, were generated using the MCMC (Table S2, Fig. 2A). Specifically, we use blocked Gibbs sampling with MCMC where we start from a random subsequence and iteratively select residues to mutate such that the final sequence follows the data distribution defined by the proteins that ESM2 was trained on and with ProtGPS and DRBERT^32^ to obtain the localization and disordered properties desired, respectively. For each protein, a plasmid was constructed that encoded the generated protein attached to an N-terminal nuclear localization sequence and a C-terminal mCherry protein. Each of the proteins were expressed in human cells together with the nucleolus marker NPM1-GFP or the nuclear speckle marker SRSF2-GFP and cells expressing both a test protein (mCherry) and the condensate marker (GFP) were isolated using flow cytometry.

Imaging of cells revealed that all ten proteins designed to assemble into nucleoli (NUC1-10) did indeed concentrate in nucleoli (Fig. 2B-D, S2). Imaging of cells expressing the ten proteins designed by MCMC to assemble into speckles revealed that six of ten (SPL1, 5, 7, 8,10) were enriched in nuclear speckles and two of ten were enriched in cytoplasmic bodies (Fig. 2D). The partition ratios of the SPL proteins in speckles were smaller than those observed for NUC proteins in nucleoli, which may be a consequence of the known distribution of speckle proteins into both nuclear speckles and much smaller RNA splicing condensates at nascent transcripts^33^. Two of the SPL proteins (SPL2 and SPL3) formed cytoplasmic foci that recruited at least one key speckle protein to the body (Fig. 2D, S4), suggesting that SPL2 and SPL3 have chemical features that assemble speckle proteins outside of the nucleus. We conclude that most proteins designed by MCMC to assemble into the two condensate compartments studied here - nucleoli and nuclear speckles – have indeed gained features that promote their concentration into these compartments.

We next conducted a sensitivity analysis for the generative process. In the multistep optimization process for each generated protein, we might expect that continuous improvement in the score computed during the optimization process should reflect the ability to generate proteins with improved compartmentalization phenotypes. As a test of this prediction, we investigated nucleolar partitioning of proteins generated at different steps during the optimization trajectory for NUC1 and NUC6 (Fig. 2E, Fig. S5A-C). Random sequences appended to mCherry, those at step 0, did not show nucleolar compartmentalization. Greater scores produced precursors to NUC1 and NUC6 proteins that tended to show improved nucleolar compartmentalization, although improvement was not continuous (Fig. 2E, Fig. S5A-C). These results suggest sampling for greater periods of time will tend to increase the likelihood of generating protein sequences with desired properties.

### Pathogenic mutations can alter protein codes

Mutations can create pathogenic effects by altering a protein’s function or altering a protein’s subcellular distribution. Because ProtGPS can accurately predict the subcellular localization of normal proteins, it might be able to identify pathogenic mutations that cause a change in the subcellular location of a mutant protein. To test this possibility, we turned to the ClinVar^34^ database, a public archive of a vast number of human variations classified for diseases. Data was collected for 205,182 mutations and ProtGPS was used to predict if the changes in amino acid sequences alter the subcellular distribution of the mutant proteins (Fig. 3A).

**Fig 3.**
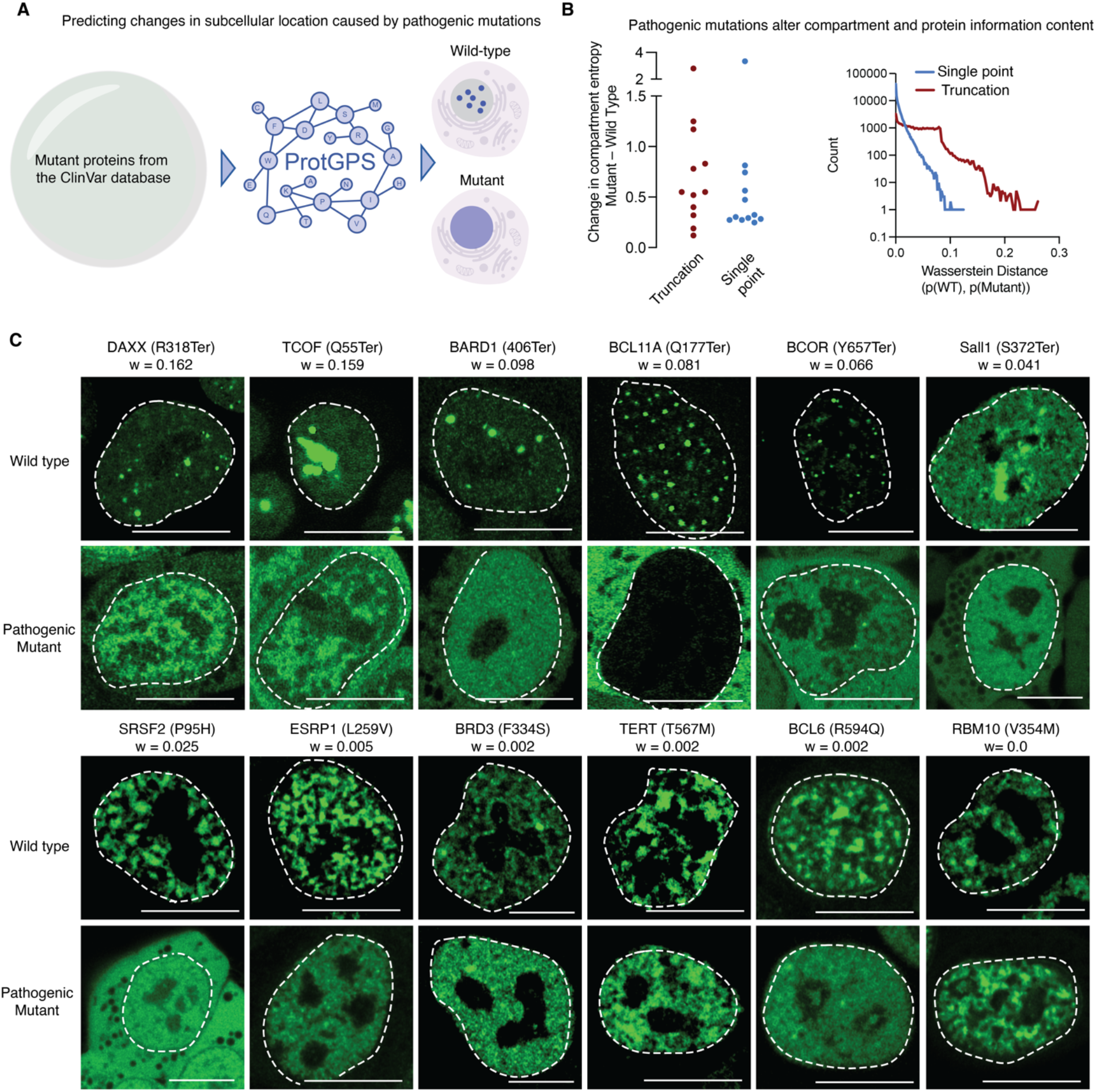
Pathogenic mutations are predicted to alter protein compartmentalization. **A**. Schematic of information flow, pathogenic ClinVar mutants caused by single point or truncation mutations were classified with ProtGPS to determine if the detected protein code was changed in the pathogenic variant. **B**. (*Left*) Dot plot showing the Shannon entropy change in compartment prediction due to single point or truncation mutation. (*Right*) Histogram showing the Wasserstein distance between the wild-type and mutant protein compartment probabilities. **C**. Live cell images of mESCs ectopically expressing wild type and truncated pathogenic variants fused to GFP, Wasserstein distance is given for each mutant as w, scale 10 microns.

We began our analysis by considering how mutations might influence the information content of condensate compartments. We computed the change in Shannon entropy^35^ of the twelve condensate compartments to learn whether the predicted information content was altered by mutation. We conducted this analysis separately for the truncation mutations (83,211), which we assumed would have major effects, from the single point mutations (121,971), which we assumed would have much smaller effects. We find that the compartment entropy is consistently higher with mutant proteins compared to the normal proteins across all compartments, with truncations producing larger effects than point mutations (Fig. 3B). Furthermore, we find that pathogenic truncation and single point mutations, when compared to normal proteins, tend to increase the Wasserstein distance^36^, a metric of dissimilarity between two probability distributions (Fig. 3B). These measures indicate that within this collection of pathogenic proteins, sequence variation may alter the predicted compartments of proteins in ProtGPS, suggesting that some mutant proteins may no longer partition selectively into compartments in the same manner as their normal counterparts.

To experimentally test the prediction that some pathogenic variants cause a change in subcellular localization, we selected for study 20 pathogenic mutations (10 truncation and 10 single point mutations) in proteins involved in a broad range of biological functions and diseases, whose normal cellular compartmentalization was well-known, and that scored across the range of Wasserstein distances (0.162-0.000) (Table S4). We then generated a panel of mouse embryonic stem cell (V6.5) lines stably expressing each protein from a doxycycline-inducible expression cassette, treated cells with doxycycline and conducted live cell confocal microscopy analysis. Differences in the subcellular localization between normal and mutant proteins would appear as changes in the fluorescence patterns displayed in micrographs. We noted that signals for all the normal proteins occurred in the subcellular locations where they are known to reside. When comparing images of normal proteins with their mutant counterparts, we found striking differences in compartment appearance for almost all truncation mutation proteins, and less striking but clear differences in compartment appearance for point mutation proteins, except for RBM10(V354M)], which scored with a Wasserstein distance of zero (Fig. 3C, Fig. S6, Table S4). Thus, it appeared that proteins calculated to have a large Wasserstein Distance tended to exhibit more dramatic changes in compartment appearance, although this relationship was imperfect. The effects of truncation mutations on nuclear localization sequences could not account for these results (Table S4). These results support the notion that ProtGPS can detect changes in protein codes due to pathogenic mutations that are demonstrable in an experimental setting.

## Discussion

Our studies suggest that proteins have evolved to harbor at least two types of codes, one for folding and another for intracellular compartmentalization. Deep-learning algorithms such as AlphaFold2, RoseTTAFold, Chroma, EvoDiff, ESM2, and others can predict a protein’s 3D structure from its linear amino acid sequence^22,37-42^. We here describe ProtGPS, which can predict a protein’s selective assembly into specific condensate compartments in cells and be used to guide generation of novel protein sequences whose cellular compartmentalization could largely be experimentally validated. The complexity of the underlying physiochemical rules for both protein folding and protein localization have proven difficult to parse using human interpretable approaches, and these deep-learning approaches therefore provide valuable predictive and analytical tools for the study of protein structure and function.

Previous studies of protein compartmentalization have suggested amino acid codes exist for some compartments. For the membrane-bound nucleus, for example, there are well-known nuclear localization sequences that facilitate the transport of protein from the cytoplasm to the nucleus^43-45^. More recently, models were used to identify patterns in protein sequences associated with specific compartments, especially those bounded by a membrane, but these did not sample a broad range of compartments and lacked generative experiments^46-48^. For nonmembrane compartments, here called condensates, there is recent evidence of patterned amino acid sequence features that can engender selective assembly of certain proteins into transcriptional and nucleolar condensates^49-51^. These observations are consistent with the concept of a protein code in amino acid sequences that promotes the selective distribution of proteins into specific compartments. Furthermore, there is recent evidence of distinctive chemical environments within condensates, suggesting that these compartments have different solvent properties ^16,51,52^. Thus, the patterns of amino acid sequences in proteins would be expected to both promote specific folding behaviors and to favor residence in compartments compatible with their solvent properties.

The patterns of amino acid sequences that occur in proteins appear overall to be highly constrained in biology^53-55^, and we suggest that this is due, in part, to the requirements for both proper folding and subcellular distribution. In our efforts to develop ProtGPS as a guide for generating novel protein sequences that promote selective subcellular distribution, we found that protein sequences sampled from collections of natural proteins were far more successful at concentrating in the desired compartment than those generated without this consideration. Analogous to the medicinal chemist’s aspiration to increase drug-like attributes such as on-target specificity and low off-target effects when developing small molecule therapeutics, designing proteins to preferentially distribute in biochemically relevant regions of the targeted cell population might improve upon their therapeutic properties^16,52,56^. In addition, exploring the chemical space of proteins naturally present in specific biological compartments may provide an especially valuable guide to the generation of optimal chemical matter directed to target proteins in specific compartments. Indeed, there are widely used and efficacious anti-cancer therapeutics that concentrate in transcriptional condensates at oncogenes^56^ due to the chemical environment of those compartments^16,52^. It is evident that similar considerations will apply to the design of protein therapeutics. We suggest that further understanding of the chemical environment established by amino acid patterns in proteins will lead to more efficacious disease therapeutics.

We conclude that ProtGPS can predict a protein’s selective assembly into specific condensates and guide generation of novel protein sequences whose cellular compartmentalization can be experimentally validated. We anticipate that future studies will advance this field by improving compartment annotation, conducting additional tests of generated proteins, deploying alternative machine learning approaches, and further exploring the effects of pathogenic mutations.

## Acknowledgments

Support was provided by NIH grant nos. GM144283 (R.A.Y.) and CA155258 (R.A.Y.), NSF grant no. PHY2044895 (R.A.Y.), Damon Runyon Cancer Research Foundation Fellowship grant no. 2458-22 (H.R.K.), and the MIT Jameel Clinic for Machine Learning in Health (R.B., P.G.M., I.C., I.M.) and Eric and Wendy Schmidt Center, Broad Institute (P.G.M., I.C.). We thank Christina Lilliehook, Alessandra Dall’Agnese, Mike Gallagher, Yana Petri, Jinyi Yang, Shannon Moreno, and Jeremy Wohlwend for helpful comments and thank Caitlin Rausch and Warbler Creative for graphical artwork.

## Author Contributions

Conceptualization: H.K., R.Y.

Methodology: H.K., I.C., P.G.M, I.M.

Investigation: H.K., I.C., P.G.M, I.M., C.V.D

Visualization: SFB, MJM, JLS, EH

Funding acquisition: R.B, R.Y.,

Project administration: H.K., R.B., R.Y.

Supervision: H.K., R.B., R.Y.

Writing – original draft: H.K., I.C., P.G.M, I.M., R.Y.

Writing – review & editing:

## Data and materials availability

Code and model weights used in this analysis are available at https://github.com/hrkilgore/protein_codes.git. Data used in this analysis are available at FigShare (DOI: 10.6084/m9.figshare.25726581).

## Supplementary materials

Materials and Methods

Figs. S1 to S6

Tables S1 to S4

Supplemental references 1-22

