## Supporting information for "Protein codes promote selective subcellular compartmentalization"

Materials and methods

*Plasmid design and purification*

Gene fragments were codon optimized for humans and purchased from Integrated DNA Technologies (IDT). Gene fragments were assembled into destination plasmids using the NEBuilder® HiFi DNA assembly kit with a molar ratio of 3:1 insert:back bone DNA (New England Biolabs, E5520S). Double stranded DNA products for CRISPR/Cas9 guide RNAs were purchased as monomer oligos from IDT, annealed in 10 mM Tris, 1 mM EDTA, 50 mM salt, heated to 95°C, and allowed to cool until reaching 25°C, over 25 minutes. Assembled duplexes were then ligated using the Quick Ligation™ Kit (New England Biolabs, M2200S) following the standard protocol provided with the product. Chemically competent E. coli cells were allowed to incubate on ice for 20 minutes with plasmids, heat shocked at 45°C for 30 seconds, before recovering on ice for 5 minutes. The transformed bacteria were then allowed to recover or SOC outgrowth medium (New England Biolabs, B9020S) for 1-2 hours, diluted 1:10, and 50 uL was spread over a 2% agar plate containing the appropriate antibiotic selection marker (Ampicillin, 100 ug/mL).

Transformed bacteria on antibiotic selection plates were then allowed to incubate overnight at 37°C. Single colonies of bacteria appearing on antibiotic selection plates were used to inoculate 5 mL of LB media (RECIPE) with ampicillin (100 ug/mL) to create an overnight culture. Overnight cultures were allowed to incubate at 37°C for 16 hours, cultures were pelleted immediately, and plasmid DNA was purified using a PureLink™ MiniPrep Kit (Invitrogen, K210011). Isolated plasmids were sequenced using whole plasmid sequencing. Restriction digests were performed to generate DNA backbones appropriate for ligation chemistry or gibson assembly reactions. Backbone plasmids were digested using restriction enzymes AfeI, BsrGI, SpeI (New England Biolabs, R0652L, R3575L, R3133L). Digest products were isolated using DNA gel electrophoresis, using a 120 V potential over 60 minutes as supplied by a Thermo scientific EC300 XL. Agarose gels were created with a 1% solution of SeaKem LEAgarose (Lonza 50004) in Tris-Acetate-EDTA buffer (Millipore-Sigma, T9650) with the addition of ethidium bromide solution 10 mg/mL, to 1 part per 20,000 (Millipore-Sigma, E1510). Gels were imaged using a Biored-Chemidoc XRST, and bands were excised while wearing UV ray eye protection. Relevant bands were isolated from gels after each run and then extracted with a razor blade from the larger gel. Gel chunks containing desired DNA bands were carefully weighed and extracted using a Monarch DNA gel extraction kit according to the manufacturer's specifications (New England Biolabs, T1020L). An estimate of DNA concentration was collected from an absorbance reading for double stranded DNA using a Nanodrop one$^{C}$ (Thermo scientific, ND-ONEC-W) and product was stored at -20°C.

*Protein design*

Proteins designed in our assays consisted of 3 (NLS-mCherry) or 4 components (all other protein sequences). These components were arranged in the following order: SV-40 NLS signal ('sequence'), an aqueously soluble and flexible linker ('SGSGSG'), a generated protein fragment, and an mCherry protein (see Table S1 and S2 for corresponding sequences).

*Tumor cell tissue culture*

Human colorectal cancer cells (HCT-116 American Tissue Culture Catalog CCl-247TM), were cultured in sterile 10 or 15 cm plates with 15 or 35 mL of DMEM (Gibco, 11965084) media supplemented with 10 % Fetal bovine serum (FBS) (Sigma F2442) and 100 units/mL penicillin (Life Technologies, 15140122), and 100 µg/mL streptomycin (Life Technologies, 15140122). Cells were cultured at 37 °C and 5 % v/v CO2 in a humidified cell culture incubator and passaged at 75 % confluency. Cells were counted to determine seeding density using a CountessTM II automated cell counter, employing trypan blue and disposable countess chamber slides according to manufacturer recommendations. Cells were tested regularly for mycoplasma using the MycoAlert Mycoplasma Detection Kit (Lonza LT07-218) and found to yield negative results. HCT-116 cells expressing NPM1-, and SRSF2-meGFP from the endogenous gene locus were previously reported.

*Stem cell tissue culture*

In these studies, we employed V6.5 mouse embryonic stem cells, a kind gift from R. Jaenisch. These cells were authenticated by short tandem repeat (STR) analysis compared to commercially acquired cells with the same name.

Cells were passaged every 1–2 days by dissociation using TrypLE Express (Gibco, catalog no. 12604), the dissociation reaction was quenched using serum/LIF medium. Stem cells were cultured in 2i/leukemia inhibitor factor (LIF) medium on tissue culture-treated plates coated with 0.2% gelatin (Sigma, catalog no. G1890) in a humidified incubator at 37 °C and 5% v/v CO2. Cultured cell lines were tested for mycoplasma regularly using the MycoAlert Mycoplasma Detection Kit (Lonza, catalog no. LT07-218) and found to yield negative results.

The composition of 2i/LIF medium is defined as 3 μM CHIR99021 (Stemgent, catalog no. 04-0004), 1 μM PD0325901 (Stemgent, catalog no. 04-0006) and 1,000 U ml−1 LIF (ESGRO, catalog no. ESG1107) in N2B27 medium.

In these experiments N2B27 medium was defined as follows: DMEM/F12 (Gibco, catalog no. 11320) supplemented with 0.5-fold N2 supplement (Gibco, catalog no. 17502), 0.5-fold B27 supplement (Gibco, catalog no. 17504), 2 mM L-glutamine (Gibco, catalog no. 25030), onefold MEM nonessential amino acids (Gibco, catalog no. 11140), 100 U ml−1 penicillin-streptomycin (Gibco, catalog no. 15140) and 0.1 mM 2-mercaptoethanol (Sigma, catalog no. m7522).

Preparation of Serum/LIF medium used KnockOut DMEM (Gibco, catalog no. 10829) supplemented with 15% FBS (Sigma, catalog no. F4135), 2 mM L-glutamine (Gibco, catalog no. 25030), onefold MEM nonessential amino acids, 100 U ml−1 penicillin-streptomycin, 100 μM 2-mercaptoethanol (Sigma, catalog no. M7522) and 1,000 U ml−1 LIF (ESGRO, catalog no. ESG1107).

*Doxycycline inducible line generation*

Piggybac lines were generated using the Super Piggybac Transposase Expression Vector (Systems Bioscience, PB210PA-1), in conjunction with protein x-lone expression cassettes (REFs for xlone). These reagents transposed proteins engineered in this study under a TR3GS doxycycline inducible promoter system. Plasmids were combined with Lipofectamine3000 reagent and Optimem media (Invitrogen, L3000015) and added to cells plated the day before at 50,000 cells/mL in DMEM (Gibco, 11965084) supplemented with 10% FBS in either 6-well or 10cm plates in accordance with manufacturer specifications. After 24 hours, their media was changed to DMEM supplemented with 10% FBS, 100 units/mL of penicillin, 100 units/mL streptomycin, and 1000-2000 ng/uL doxycycline halcylate (Millipore-Sigam, D9891) in water.

Twenty-four hours after induction with doxycycline, cells were prepared for sorting by washing cells twice with 10 mL of phosphate buffered saline prior to the addition of 1.5-3 mL of TrypLE to trypsinize the adherent cells for 5-10 minutes at 37 °C. The trypsin reaction was then quenched by the addition of 5 mL of DMEM (Gibco, 11965084) containing 10% FBS, 100 ug/mL of penicillin and streptomycin (Life Technologies, 15140122). Cells were pelleted in 15 mL conical vials at 500 RPM using a table top centrifuge, and resuspended in Dulbecco's phosphate buffered saline containing magnesium and calcium (GibCo 14040117), and filtered into a 5 mL polystyrene round-bottom tube outfit with a cell straining cap (Corning, 352235).

Cells were sorted by flow cytometry as described below for double positives colon cancer (HCT-116) cells expressing green fluorescent protein tagged SRSF2 or NPM1 from the endogenous locus and the target mCherry protein (see Table S1 and S2 for sequences). A homogenous population of cells expressing only SRSF2-meGFP or NPM1-meGFP from the endogenous locus was used as a positive control for meGFP expression and a negative control for mCherry expression. Double positives were collected into 1.5 mL Eppendorf tubes containing 500 uL DMEM containing 10% FBS, 100 ug/mL of penicillin and streptomycin and stored on ice until they could be transferred into 12-well dishes. Sorted cells were cultured for 7 days or until approaching confluency in 12-well dishes.

At approximately 75 % confluency, cells were taken up into solution following the protocol for TrypLE and washes given above, the concentration of cells was established using a CountessTM II automated cell counter, employing trypan blue and disposable countess chamber slides according to manufacturer recommendations. Each population of population of double positive cells was then diluted to 0.85 cells / 100 uL in DMEM containing 10% FBS, 100 ug/mL of penicillin and streptomycin. A multichannel pipette was then used to transfer 100 uL of the diluted cell solution into four 96-well plates and cells were allowed to grow for 7-14 days until single colonies were identified, with media changes occurring on days, 4, 8, and 11. Clonal cell populations were replated in 96-well imaging plates and imaged with confocal microscopy to identify those clonal populations possessing the desired double positive phenotype. Chosen clonal populations were then transferred into 12 wells and allowed to grow to confluency before replating in 10 cm dishes for analysis.

Production of stable mouse embryonic stem cell lines was performed by cloning WT and mutant gene sequences using NEBuilder HiFi DNA Assembly (NEB) into a doxycycline-inducible, N-terminal mEGFP-tagged expression construct with a hygromycin-resistance gene (pbfh-GFP), which was integrated into mESCs using the PiggyBac transposon system (Systems Biosciences). 0.5 x $10^6$ wildtype mESCs were plated in 6-well format and simultaneously transfected with 1 µg of the expression vector and 1 µg of the PiggyBac transposase using Lipofectamine 3000 (ThermoFisher), according to manufacturer instructions in serum/LIF media. The next day, media was changed to 2i, and cells were split into 100 mm gelatin-coated plates with 2i-media supplemented with 500 µg/mL hygromycin (ThermoFisher) for selection. Selection media was exchanged every day and un-transfected control cells were monitored to assess selection.

*Flow cytometry*

Samples were sorted using a BD FACS Aria. Green fluorescent protein and mCherry signal was used to identify colon cancer (HCT-116) that were expressing NPM1 and SRSF2 meGFP fusion proteins in addition to the mCherry proteins incorporated as described in the section "Doxycycline inducible line generation." Double positive cells were collected when both channels had relative signal 10-fold above the background signal produced in the absence of mCherry and within the region defined by the signal found in the meGFP-SRSF2 and meGFP-NPM1 control cell lines. Double positive lines were sorted into 1.5 mL Eppendorf tubes containing in 1 mL of media and stored on ice until plating.

*Live cell imaging*

Endogenously tagged HCT-116 cells expressing NPM1-meGFP or SRF2-meGFP and proteins from Table S1 and Table S2 were seeded at 50,000 cells/mL on an imaging plate. Imaging plates used were sterile Cellvis 96-well glass (Cellvis, P96-1.5H-N) bottom plates with #1.5 high performance cover glass (0.17 ± 0.005 mm), or sterile Cellvis 384-well (Cellvis, P384-1.5H-N) glass bottom plates with #1.5 high performance cover glass (0.17 ± 0.005 mm). Cells were plated 48 hours prior to the experiment in DMEM containing 10% FBS, 100 ug/mL of penicillin and streptomycin. Twenty-four hours before imaging, cell media was changed to DMEM containing 2000 ng/uL doxycycline halcylate in water (Millipore-Sigma, D9891) 10% FBS, 100 ug/mL of penicillin and streptomycin. Cells were maintained at 37 °C with 5 % v/v $CO_2$ in a humidified chamber over the course of the imaging experiment.

*Imaging instrumentation*

Live cell confocal micrographs were recorded with a Zeiss LSM 980 Airyscan 2 Laser Scanning confocal with a 1.4 NA ×63 Plan Apo objective and running Zeiss Zen Blue v.3.5. Cells were maintained at 37 °C and 5% v/v $CO_2$ in a humidified chamber throughout the experiment. Images were recorded using 405 nm at 25 mW, 488 nm at 25 mW, 561 nm at 25 mW or 639 nm at 25 mW diode lasers as required.

*Imaging data analysis*

Our image analysis approach was designed to compute the partition ratio of nucleolus and splicing speckle targeted proteins within the surrounding spatial region in each cell. Regions were defined using Zeiss Zen Blue image analysis software. Nucleophosmin (NPM1) is a scaffold and marker of the granular cluster of the nucleolus and serine arginine-rich splicing factor 2 (SRSF2) is a marker for nuclear speckles. Nucleoli and nuclear speckles were identified using the 488 nm excitation band, which could indicate the distribution of nucleophosmin (NPM1)-meGFP and SRSF2-meGFP fusion proteins in the cell. Global threshold-based detection using the following options enabled identification of the nucleolus: a three-sigma threshold approach, a minimum object area of 10 pixels$^2$ (a size constraint of 50 pixels$^2$ was used to cull erroneous calls of nuclear speckles), object expansion employed a closed binary criterion, gaussian smoothing, and signal segmentation was performed using watersheds. Regions found outside of the target condensates were identified using Otsu thresholding (light-regions), without gaussian smoothing, object expansion was set to none, and watersheds.

Average signal was computed for inside of the nucleolus ($I_{nucleolus}$) and in the nucleoplasm ($I_{nucleoplasm}$) to compute a partition ratio $K_{nucleolus} = I_{nucleolus} / I_{nucleoplasm}$. Images collected of mCherry signal using the 561 nm excitation laser were analyzed used to calculate, $K_{nucleolus}$, providing the partition ratio of mCherry proteins in regions defined above.

Average signal within splicing speckles and cytoplasmic regions defined by the accumulation of SRSF2-meGFP was computed using $I_{SRSF2.}$ Reference regions, such as the nucleoplasm were using $I_{nucleoplasm}$ or $I_{cytoplasm}$, which was manually defined in Zen blue. Images collected of mCherry signal using the 561 nm excitation laser were analyzed used to calculate, $K_{SRSF2}$, providing the partition ratio of mCherry proteins in regions defined above.

*Identification of pathogenic mutations*

Pathogenic mutations were collected from ClinVar database and annotated following the approach of Banani et al. 2022[1-7]. Variants associated with Mendelian diseases were obtained from HGMD v2020.4 [4], ClinVar [1] and in hg38. AACR Project GENIE v8.1 [3] and various TCGA [2] [8] and TARGET studies via cBioPortal were used to collect cancer variants.
(cBioPortal study
identifiers: ucec_tcga_pan_can_atlas_2018, skcm_tcga_pan_can_atlas_2018, coadread_tcga_pan_can_atlas_2018, luad_tcga_pan_can_atlas_2018, stad_tcga_pan_can_atlas_2018, lusc_tcga_pan_can_atlas_2018, blca_tcga_pan_can_atlas_2018, brca_tcga_pan_can_atlas_2018, hnsc_tcga_pan_can_atlas_2018, cesc_tcga_pan_can_atlas_2018, gbm_tcga_pan_can_atlas_2018, lihc_tcga_pan_can_atlas_2018, ov_tcga_pan_can_atlas_2018, lgg_tcga_pan_can_atlas_2018, esca_tcga_pan_can_atlas_2018, prad_tcga_pan_can_atlas_2018, paad_tcga_pan_can_atlas_2018, kirp_tcga_pan_can_atlas_2018, kirc_tcga_pan_can_atlas_2018, sarc_tcga_pan_can_atlas_2018, thca_tcga_pan_can_atlas_2018, acc_tcga_pan_can_atlas_2018, ucs_tcga_pan_can_atlas_2018, laml_tcga_pan_can_atlas_2018, dlbc_tcga_pan_can_atlas_2018, thym_tcga_pan_can_atlas_2018, meso_tcga_pan_can_atlas_2018, kich_tcga_pan_can_atlas_2018, tgct_tcga_pan_can_atlas_2018, chol_tcga_pan_can_atlas_2018, pcpg_tcga_pan_can_atlas_2018, uvm_tcga_pan_can_atlas_2018, wt_target_2018_pub, all_phase2_target_2018_pub, aml_target_2018_pub, nbl_target_2018_pub, and rt_target_2018_pub).

Liftover [9]was used to convert the genomic coordinates for different cancer variants from hg19 to hg38. We did not consider deletions larger than 100kb in this analysis. Protein coding sequences changes associated with variants in our study were mapped to the set of 20,394 human proteins using Ensemble VEP v102 and ID mappings between Ensemble and UniProt [10]. We considered the pathogenic mutations in the context of the canonical isoforms in this study, which represent the best characterized set of isoforms. Isoforms are selected from criteria such as prevalence, similarity to other homologs, and without consideration of other information (e.g., sequence length) [11]. A collection of n= 2,644,688 DNA variants (62% of all variants located within source data sets) were mapped onto the 20,394 canonical protein isoforms found within UniProt. All variant were counted as protein variants—i.e., DNA variants resulting in the same protein-coding alteration on different DNA sequences, were counted as the same. Synonymous variants were excluded from our analysis. For non-synonymous variants, only the primary and most severe protein-coding change associated with a variant was considered based on the established hierarchy of mutation effect severity conveyed by variant annotations in Ensemble.

Mendelian variant pathogenicity was classified from the designations of their clinical significance for ClinVar variants (pathogenic or likely pathogenic) or of variant class for HGMD variants (DM or DM?). Cancer variant pathogenicity was determined by assessment of variants for their inclusion in CIViC[12], their inclusion in the list of CGI's validated oncogenic mutations, or oncogenicity designation in OncoKB v2.10 (predicted oncogenic, likely oncogenic, or oncogenic) [13]. Definitions of pathogenicity rely on computation prediction of pathogenicity, but are less dependent upon computation prediction than clinical biological/functional, or evolutionary evidence of pathogenicity[14,15].

Among pathogenic mutations, we chose to investigate those that might be readily discernable as influencing structure and assembly. Nonsense and frameshift variants were considered together to be truncating variants and assessed for their predicted propensity to elicit NMD. Predictive rules for NMD were obtained from prior work [16]. A truncating variant was considered to elicit NMD if the corresponding premature stop codon it introduced occurred (i) >200 residues C-terminal to the start codon; (ii) >50 residues N-terminal to the final exon-exon junction; and (iii) in an exon ≤400 base pairs in length. Mutations were then be subsampled to include only those identified as a single point or truncation mutations, leading to 205,182 protein sequences. The resulting mutant protein sequences were then classified using ProtGPS.

*Prediction of nuclear localization and nuclear export signals*

Nuclear export and localization signals identified from UniProt *motifs* possessing a description "Nuclear localization signal." Nuclear localization and nuclear export signals within NLSdb [17] that were filtered to be comprised of features annotated as *Experimental* or *By expert*.

*Compartment Classification*

To train ProtGPS's compartment classifier module, we collected a dataset of 5,480 proteins from UNIPROT and CD-CODE [11,18] covering 12 condensates, consisting of nuclear speckles, p-bodies, PML-bodies, post synaptic densities, stress granules, chromatin, nucleoli, nuclear pore complexes, Cajal bodies, RNA granules, cell junctions, and transcriptional condensates. We randomly assigned 70% of protein sequences to training, 15% to development, and 15% to test, yielding 3,766, 785, 803 sequences in each split, respectively.

Our model utilizes only the protein sequence to obtain a binary prediction for each of the 12 condensates. Specifically, we initialize a sequence encoder using the protein language model ESM-2 with 8 million parameters (esm2_t6_8M_UR50D) [19], and utilize a 2-layer feed-forward neural network with a hidden dimension of 512 as the classifier head. The classifier is implemented with batch normalization. From the ESM-2 model, we obtain a 320-length feature vector per residue. We take the mean embedding across residues to obtain a 320 embedding of the protein sequence, and pass it to the MLP. We train the model end-to-end for 90 epochs in half precision. We use a batch size of 10, an initial learning rate of 0.001, an exponentially decaying learning rate schedule with a decay rate of 0.91, and a dropout rate of 0.1. The model is optimized with the Adam algorithm[20] with default parameters. All models are implemented in PyTorch (v2.0.0+cu117) and PyTorch Lightning (v1.6.4).

*Protein generation with ProtGPS: Autoregressive Greedy Search Generation*

Our first attempt at generating proteins possessing chemical codes for different compartments utilized a greedy search algorithm. Given the mCherry sequence, we add to the N-terminus a random subsequence of length $\ell = 150$. This sequence is then iteratively mutated at each position of the subsequence. At each step, we predict the localization of the protein when mutating the current position to all 20 possible amino acids. We keep the top 3 sequences predicted to localize to the desired compartment. For each of those 3 sequences, we repeat the process of mutating the next position (obtaining 3x20 sequences) and keeping only the top 3 scoring proteins. Once all $\ell$ positions are explored, we choose the single protein most likely to localize to the target compartment among all those generated. The full generation process is repeated 40 times to obtain 40 sequences.

*Protein generation with ProtGPS:* MCMC Generation

We adapt the framework presented in Verkuil et al. [21] to generate novel sequences that lead to the localization of mCherry to specific condensates. In particular, we aim to sample sequences $x$, where the first $\ell$ amino acids are designed computationally and the rest of the protein corresponds to the mCherry sequence. We guide the generation such that (1) the newly generated subsequence follows the natural distribution of protein sequences, (2) the subsequence is predicted to be disordered, and (3) the full protein is predicted to have the desired localization phenotype (e.g., localizing to the nucleolus). We use blocked Gibbs sampling with MCMC[21] where we start from a random subsequence, sample a backbone structure $y$, then update the sequence given the current backbone. This process generates sequences according to the data distribution defined by the proteins that ESM-2 was trained on. Doing so results in sequences that are expected to follow the distribution found in the natural world.

However, our aim is to specifically generate IDRs that are consistent with the chemical space of the condensate we are targeting. To do so, we use ProtGPS to condition the generation process on the likelihood that the full sequence (with mCherry) localizes to the desired condensate. In other words, we sample subsequences from the space of proteins that ProtGPS predicts to localize in our target condensate. To ensure the novel subsequence does not have a definite 3D fold, we use a predictor of protein disorder as a further constraint. Formally, we sample from the joint distribution

$$x, y \sim \mathrm{p}(\mathrm{x}, \mathrm{y} | \mathrm{c} = \mathrm{C}, \mathrm{d} = 1)$$

where $c$ is the condensate compartment we are targeting, and $d$ indicates whether the generated subsequence is disordered. Since the sequence can fully determine structure, the backbone structure $y_{sampled}$ is obtained as in[21]:

$$y_{sampled} \sim p(y|x)$$

However, we sample a new amino acid sequence (keeping the mCherry sequence fixed) as:

$$x' \sim p(x|y = y_{sampled}, c = C, d = 1)$$

We consider the likelihoods that a sequence localizes to a specific condensate and that it contains an IDR to be conditionally independent. So, we obtain:

$$\mathrm{p}(x|\mathrm{y}_{\mathrm{sampled}},\ c = \mathrm{C},\ d = 1) \propto p(c = C|x, y_{sampled})p(d = 1|x, y_{sampled})p(x|y_{sampled})$$

Therefore, we add two terms, $E_{condensate}$ and $E_{IDR}$, to the original energy-based MCMC sampling[21]:

$$\mathrm{E}(\mathrm{x}) = \lambda_{\mathrm{p}} \mathrm{E}_{\mathrm{projection}}(\mathrm{y} = \mathrm{Y}|\mathrm{x}) + \lambda_{\mathrm{LM}} \mathrm{E}_{\mathrm{LM}}(\mathrm{x}) + \lambda_{\mathrm{n}} \mathrm{E}_{\mathrm{ngram}}(\mathrm{x})$$

$$+ \lambda_c E_{condensate}(x) + \lambda_{IDR} E_{IDR}(x)$$

where

$$E_{condensate}(x) = -log\, p(c = k \mid x)$$

$$E_{IDR}(x) = -\sum_{i=1}^{\ell} log\ p(x\_i\ \in IDR|x)$$

Note that we use the full sequence to predict localization, but we calculate disorder only for the first $\ell$ residues, where $\ell$ is the length of the IDR we seek to generate.

As in[21], we use ESM-2 (esm2_t33_650M_UR50D)for the language model and protein structure samplers. We use ProtGPS to estimate the likelihood the generated sequence localizes correctly ($p(c = k \mid x)$), and the DR-BERT model[22] to predict the disorder of each residue ($p(x\_i\ \in IDR|x)$). We generate sequences of length 100 at the N-terminus of the mCherry protein sequence, keeping the mCherry sequence fixed throughout the process. We set the weights for each energy term as $\lambda_{\mathrm{p}} = 3,\ \lambda_{\mathrm{LM}} = 2,\ \lambda_{\mathrm{n}} = 1, \lambda_c = 1, \lambda_{IDR} = 1$. Since we do not intend for the sequence that we generate to have a highly ordered structure, we stop the generation process when the sequence has a likelihood of $p(c = k|x) > 0.85$ for 10 consecutive steps (instead of performing 170,000 MCMC steps). We use a warm-up of 1000 steps. For all other parameters, we use the default values. We use a different seed to initialize each process.

*Analysis of Predicted Localization in Pathogenic Mutations*

For each compartment, we collect the predicted scores for the WT and mutated sequence, and create a histogram for each with 100 bins in the range of 0 to 1. We normalize the histograms and get two distributions for which we compute the Shannon entropy. For each pair of WT and the corresponding mutated sequence we also calculate the Wasserstein distance between the predicted scores of the two sequences. A set of predicted scores for a given sequence can be regarded as the localization distribution for that sequence. Hence, the Wasserstein distance reflects the optimal transport cost between the two distribution – the sum of minimal changes required in compartment prediction scores to match the two distributions.

Supplemental Tables

**Table S1**. Autoregressive greedy search generated N-terminal peptides created to target mCherry to the nucleolus.

| Sequence ID | Sequence |
|---|---|
| 400 | KRIRSIRMMVKYMGEAFEYEGPCHTVGGKFTCHGICSYIHHPRPVMGGNYAWSTRSY<br>WVMMAICTVPKYFNGDICMVSDNGHGCCVGMGLACLCQKHYKMKHMNHDAHTEY<br>NHTAEWHHDWEEWFLECMANAPWMAKMEAQIDFKMKGDT |
| 403 | KKRMWDRQRFSTSFCYTGYIESIWGWGLYDAMRTVAPQKWPKVITIMWHAFASPYPK<br>LCLQAHITNCGCHMRTVFIRCPWTFSGEIKHCGWCAWCDRWLQCFAHADMNVACSYS<br>KQKPPPQRNVDRIHCTAVKPCIPRLPYVPHPGPYCC |
| 404 | AKKISCHLRILCYDRTSSMRWYAQGRAMRKAKEIRCNNHVQYSCKTIWCRMGIMIMEG<br>NCWETWITYHYTTAHPGKNWHVRRWDNMQKECNHAGWPRPRLIWHTHGLKHAMMS<br>EKSEPHYNQDEACFYNKGMYEYSMMEHHDMDFVTCVS |
| 674 | IEEEEIFMHRRRMGDWIQSKCKQCHEWNIKNHCEVWFKWRVSYCRCFNSFNCGQVRN<br>PCQMCDHLVGTTTFMQRKEKDVGEILQMRHPIGRCAVFCSHQPKHNFISHETCRAQGR<br>WMLNSEEDMEVPIACADCGAWCCIHWEEDHSWPCT |
| 440 | GIETNKYRIYGAWDWIVASIIVQGVCDFHYTTTKEALRFKYIFGKMSWKHGCAIERRCN<br>NIGAIIDKHRFHEHIRAEANNAWAWPCAMAIDPIFRCGWFWLKVRPERYVKPKKFWKH<br>KENEDDHIKINTEIHHIWNWMMCWWHDDTKKSV |
| 401 | MGVDEKLEMTALFCIGIMGLSWQCCRSEVIMCCDDDTRHIMCMVVHYSIRCSYQRMK<br>YAMRDKKYFIWKMFRSKIGHRKSICLWRLHGIHKPWQTQWTSAAKDHKPYPDKHQNY<br>EHHYYDVRNTCRKIWHPSGYHADMDEWTMNNLEDAE |
| 440 | GIETNKYRIYGAWDWIVASIIVQGVCDFHYTTTKEALRFKYIFGKMSWKHGCAIERRCN<br>NIGAIIDKHRFHEHIRAEANNAWAWPCAMAIDPIFRCGWFWLKVRPERYVKPKKFWKH<br>KENEDDHIKINTEIHHIWNWMMCWWHDDTKKSV |
| 505 | GDKNYAMMKGDNTEAWGGWKRAYSHTHHTKHCIRPRVKCWHYRIKKHSFIHGDWCC<br>YNRVIWGPEKRWWCQHFYGDENHDIKENWEEDRECAPISPGHPILREAVRRRFLRYTR<br>CNIRLSFIRMPEGTSSVQHFTDIQRVKWFDYGFPVH |
| 506 | LKHDGNHHHGCAKNVIHRYDEQRCDHTAADHINCVYGGLIKAQDTATWSHMTLTMY<br>MCHFNSHRIPTCKQWWRWYECTVPRLQPMERKDITPRHRGRRLGGKTMYPEWMNRC<br>WNHTNGINHECSGTWDDRSHHGGCPYHCDDGIRNECTS |

**Table S2.** Markov Chain Monte Carlo generated N-terminal peptides created to target mCherry to the nucleolus (NUCX) or nuclear speckles (SPLX).

| Name | Sequence |
|---|---|
| **NUC1** | FMLVSTLWWKQKRLNNAVRTHTKFLTTINNPWRDFCSHRKKYCQKRKHEHATLKSWGTNN GSRRAAGICSGYGPEHSPDANTVKHCCIDYDSIDPIRCTR |
| **NUC2** | HFMRIADRKVMHHGCAKQGNSWNHIGQKPCCSKVKKGEQSQKADAVVWGVKCHMKWE ARSQCNQSFEKMQLHCPMSCRVQESSHNQHNIQPKANHQAMIH |
| **NUC3** | ATDYRQEGLKMETQMSVTDAMIPSGPVKWGCNPNSKSKQKPTSVRQATHGTAWTQESHVW NIWGIPCQLHADTHADPFEWKGVAHTADPVNHDRANRNES |
| **NUC4** | DWKWRMYGEWDSTGTVMGEWGHRHCDTVQAICWVNRLYRKKEDPQAKHDFRAHQLPM AQNKPKQHQCKKEEGILEPSKGVGGKGIRMWWDPEYRIYQEPL |
| **NUC5** | GRPFRRFKKWEELDGPPIGEQLQGRLRETAYPLKEKIHTHHIFGRMVKTDWLPCWQHSGHLI CRRMWSIFPEPTLKKKMDGHSNPHGAEGSQHKDFDPWS |
| **NUC6** | NCLETANAEMDEPHDKILHEPRKAVRYQHHGQEYDRLQWPVTHPTFAESEMEKQRHYVHC DRKRWKKCRIEEEKRQKLPHEPLEVSPVKHCPFEAEEYNG |
| **NUC7** | HGQNRRRKNIGTLKMHTIRGFFPMFSEIRNNHTFTIHGSKSFNSDFQDQNLHCHDRMMHLQI SDSMNNTGEEWMTEKVNSLPRKGKSGGPPYKPKVWSVQ |
| **NUC8** | FMDDVLWQLHARQSFRYAHHFPGPVNSRKHFTHTICSDVDKNTRMGEDNMVPMCMPEAEY ICPIDDLSLARSHKQRDMSTIFQETNLKVSNKQWRRPWLQ |
| **NUC9** | HNVHRMNKSKLSLTLKRQPTITAMHFEASVSNHWKGFPSSNVVAHSGYYKEIAPHVTEQAN MDQGVGMGIRSQSHTSQLNEIDNEPPGEAKESSAGCASY |
| **NUC10** | LSPTWCDEVANDQQPTGNQAETHICNSIKGQSEAMGEHNNMQHVAGDWEKYMEPVYPHGE EMDNPYAMCDGHFCPEIIKSLGGRANNKTQNGQYQINKHT |
| **SPL1** | PGVPHQTLRHHIRPHEHIHAFKDRNWEGKTKGPEFNEYHNAEFHHHGTNESHCSERAKFRF HQQRQTPHREIIESLSEEWNSTQKECHHRTTKEFVKCGK |
| **SPL2** | VNDITDVEMAVGRVPREGGNATERCYACFHHLDDYDLHQQMHGRDAPHMRNNSYKKAAH SEHINEVDHQGLQSDVEEYEGVMNEDTFKYMADERDCSPRN |
| **SPL3** | TKIKKHRSTPNMIQSPVTYPDEDHTNNHAGWKTTKAAAPKFRCAARQINRTAMMRCENFAI TIDDMPSQDWPHKDDHGAGDDKKDCMPARYDGHTEETND |
| **SPL4** | FFDDVLWQLHARQSFRYAHKFPGPVNERKHFTHSRCSDVDKNTRMGEQNKVPMCMPEAEY ICPIDDLSLARSHAQRDMSTSFCETYPKVSNTQWRRPHLQ |
| **SPL5** | EKSHMHGLSMHNCHCGGMSCHHYQQPKMHAVSYKKFVNYGPVEDTLGARDEFVYHVRRS EKRREMNNFEPWQFHTKTKTRHHKQSSHEGTWKWPAPQFHP |
| **SPL6** | RRFRASIRLVHACGHNHEGKRPFGERWPCEDDKHKPMENQLMKCPFSLMHQQMAYMMEM GDEWHPTMHYHTHMHAPMAEETYKTKVYNSYYGLGWWVDPM |
| **SPL7** | THDEYSYHTRNTRNGFAFDRKDTGRSWGEYNQFKQTGADVNTDTRPLHRPAPKNNTRLYA GRGLSRTKCKLERTTSRHQERHTGNNPEFASNCVSEPAFP |
| **SPL8** | CEYLHARTFSTRVPHAAISTVSPSKDYEDNGYHPAADDCPADSHCYPTMYDKTQWHEYRWH DTQHPSIDQKGNVSAHSEFHQHTGCNPAFFSKALNVMQY |
| **SPL9** | YEFFFPERLVRISQAPKLKELEGTGMRKEPPSTKCTMCFNDLCMLVLHGRIWRIKQQDVKNN PSDAMKVTEGENAADKHDHRKGSRHPMYCCPMCDCDFM |
| **SPL10** | NTSTGVEKHKRVTNQKRDDCTKSCCMISQKIAVARDGHDEVTAPPYTRYTHDVPCYGQTSV HKPRLNFKTADVMECDLSGHCSFEKKIDKETQNDEKMLD |

**Table S3.** Statistical analysis of data reported in Fig. 2d.

| Name | # of cells assayed | p-value (KS-test) |
|---|---|---|
| **NUC1** | 56 | < 0.0001 |
| **NUC2** | 32 | < 0.0001 |
| **NUC3** | 33 | < 0.0001 |
| **NUC4** | 34 | < 0.0001 |
| **NUC5** | 33 | < 0.0001 |
| **NUC6** | 51 | < 0.0001 |
| **NUC7** | 16 | < 0.0001 |
| **NUC8** | 18 | < 0.0001 |
| **NUC9** | 31 | < 0.0001 |
| **NUC10** | 15 | < 0.0001 |
| **SPL1** | 10 | 0.0193 |
| **SPL2** | 13 | < 0.0001 |
| **SPL3** | 15 | < 0.0001 |
| **SPL4** | 12 | 0.7102 |
| **SPL5** | 16 | < 0.0001 |
| **SPL6** | 17 | 0.4129 |
| **SPL7** | 17 | 0.0013 |
| **SPL8** | 14 | 0.2343 |
| **SPL9** | 15 | 0.2923 |
| **SPL10** | 13 | 0.9242 |

**Table S4**. Table of pathogenic variants considered experimentally in this study.

| Protein | Mutation | Magnitude of effect on distribution | Wasserstein Distance (WT, Mutant) | Fraction of predicted NLS and NES signals mutated |
|---|---|---|---|---|
| **DAXX** | R318Ter | Major | 0.162 | NES 3/6<br>NLS 11/14 |
| **TCOF** | Q55Ter | Major | 0.159 | NES 2/4<br>NLS 50/53 |
| **BARD1** | R406Ter | Major | 0.098 | NES 1/4<br>NLS 2/6 |
| **BCL11A** | Q177Ter | Major | 0.081 | NES 2/4<br>NLS 1/3 |
| **BCOR** | Y657Ter | Major | 0.066 | NES 3/4<br>NLS 13/13 |
| **SALL1** | S372Ter | Major | 0.041 | NES 3/5<br>NLS |
| **SRSF2** | P95H, S54H | Major, Minor | 0.025, 0.003 | NES 0/0, 0/0<br>NLS 0/5, 0/5 |
| **ESRP1** | L259V | Minor | 0.005 | NES 0/6<br>NLS 0/0 |
| **BRD3** | F334S | Major | 0.002 | NLS 0/24 |
| **TERT** | T567M | Minor | 0.002 | NES 0/3<br>NLS 0/5 |
| **BCL6** | R594Q | Minor | 0.002 | NES 0/3<br>NLS 0/1 |
| **RBM10** | V354M | Major | 0.000 | NES 0/3<br>NLS 0/17 |
| **MECP2** | R186Ter | Major | 0.067 | NES 1/3<br>NLS 11/16 |
| **DYRK1A** | Q547Ter | Minor | 0.065 | NES 0/3<br>NLS 0/8 |
| **ASXL1** | R693Ter | Minor | 0.023 | NES 1/2<br>NLS 0/13 |
| **BRCA1** | D720Ter | Minor | 0.020 | NES 0/2<br>NLS 0/2 |
| **ENC1** | P404Q | Minor | 0.001 | NES 0/3<br>NLS 0/2 |
| **CBX5/HP1a** | V21L, W142C | Minor, Major | 8.33E-05,<br>4.16E-04 | NES 0/1, 0/1<br>NLS 0/6, 0/6 |

Supplemental Figures

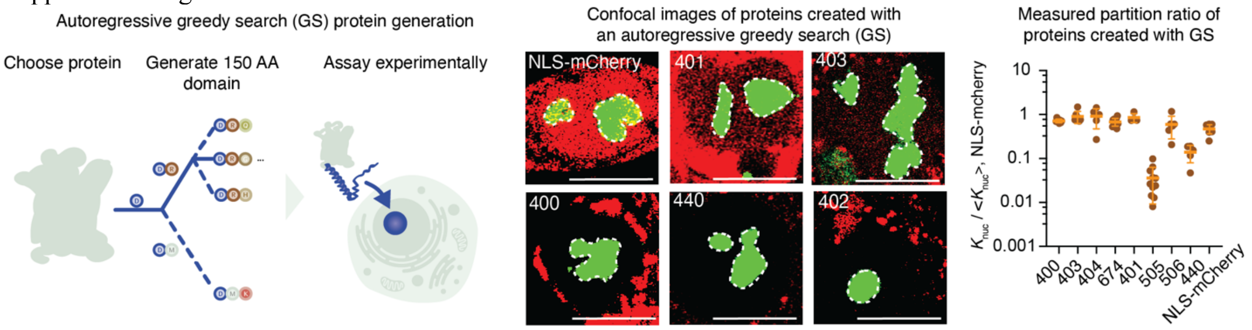


**Fig. S1. Protein generation using an autoregressive greedy search algorithm.** (*Left*) Schematic showing the approach to generating proteins using an autoregressive greedy search algorithm guided by ProtGPS. The nucleolus is shown in green (indicated by NPM1-GFP) proteins were generated to target the nucleolus. (*middle*) Confocal micrographs of GS proteins targeted to the nucleolus expressed in colon cancer (HCT-116) cells tagged at the endogenous locus of nucleophosmin (NPM1) with green fluorescent protein (GFP) to indicate the nucleolus (488 nm excitation, green, 561 nm excitation red, overlap, yellow). Dashed lines indicate the perimeter of the nucleolus. scale: 10 microns. (*Right*) Dot plot on a log scale showing the partition ratios of GS proteins in the nucleolus relative to the nucleoplasm ($K = I_{nucleolus} / I_{nucleoplasm}$).

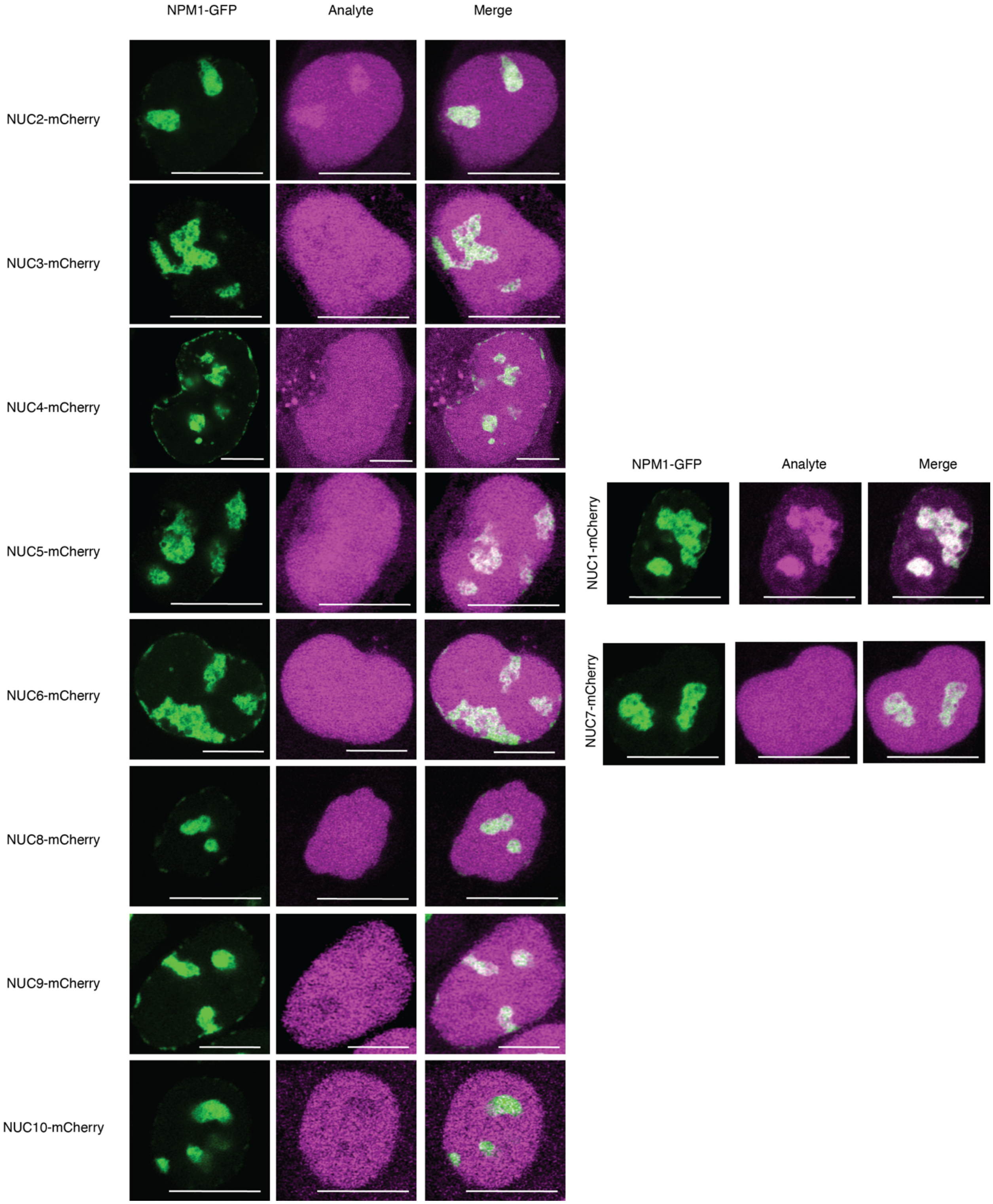


**Fig. S2. Live cell confocal microscopy images of NCMC generated proteins targeted to nucleolus.** Live cell images of colon cancer (HCT-116) cells expressing NPM1-GFP (Green) from the endogenous NPM1 locus and induced expression of the indicated nucleolus targeted protein, scale 10 microns. Some images are reproduced here from main text Fig. 2C.

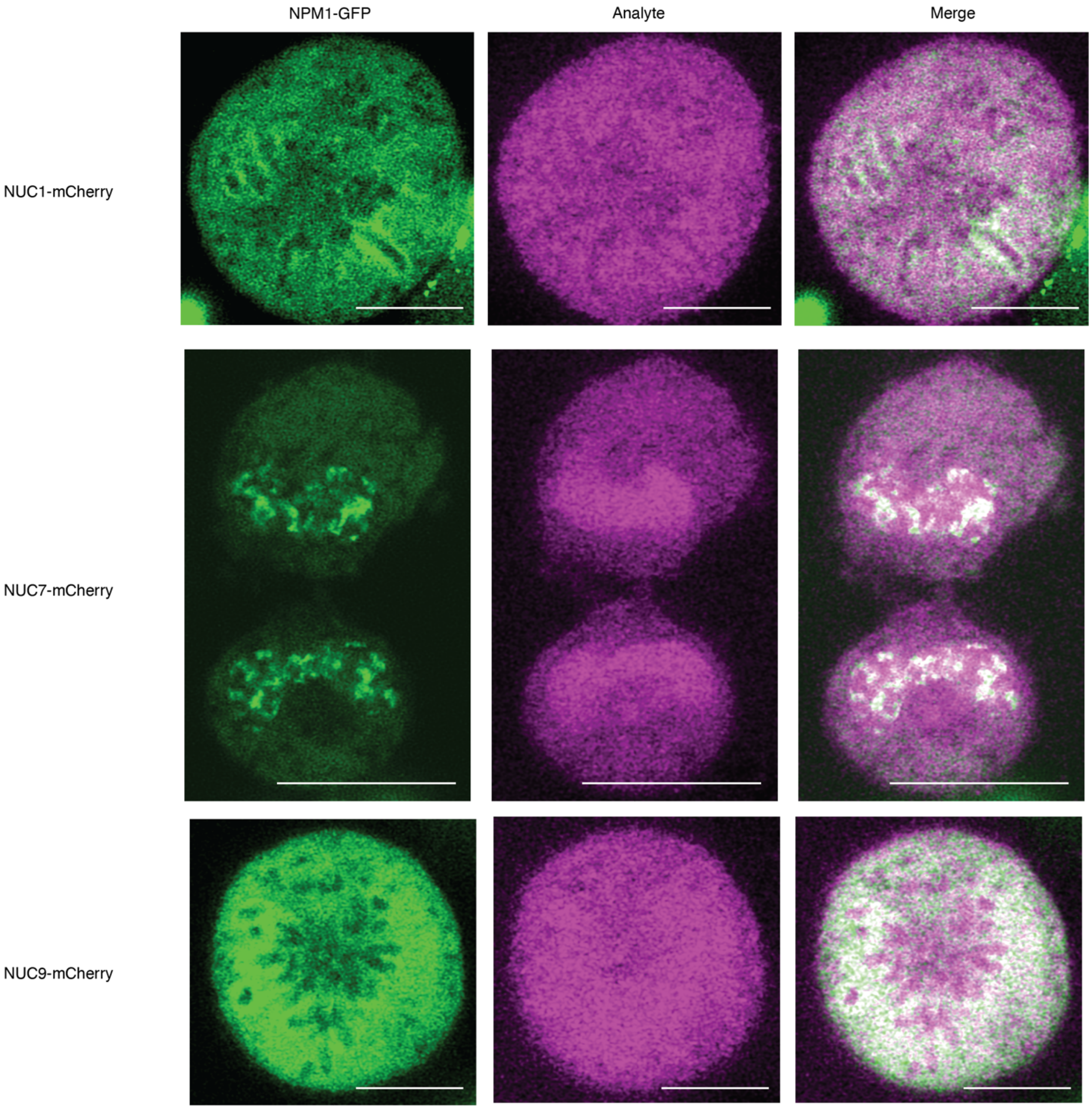


**Fig. S3. Confocal microscopy of some nucleolus targeted proteins at different stages of the cell cycle.** Live cell images of colon cancer (HCT-116) cells expressing NPM1-GFP (Green) from the endogenous NPM1 locus and induced expression of the indicated nucleolus targeted protein, scale 10 microns.

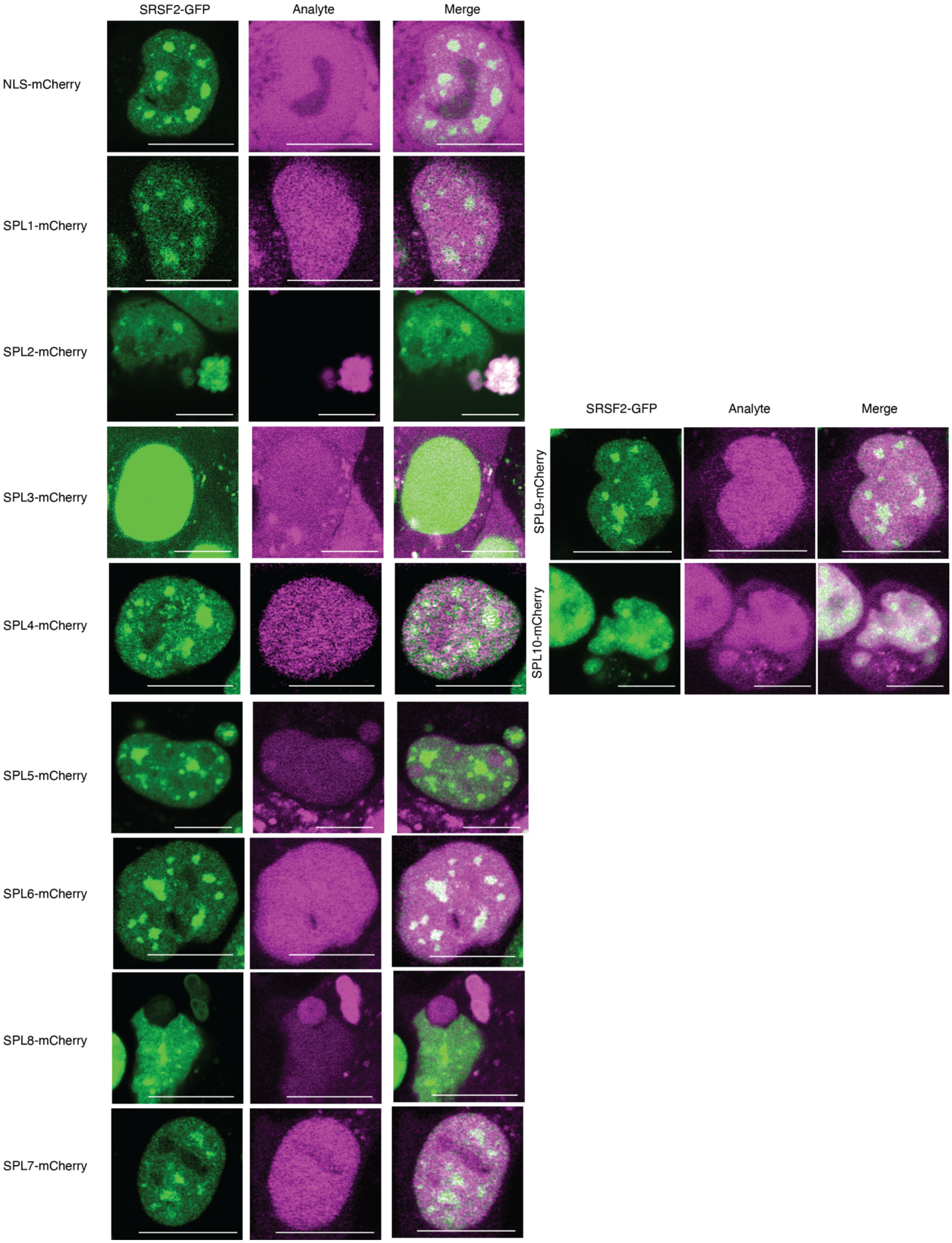

SRSF2-GFP
Analyte
Merge
NLS-mCherry
SPL1-mCherry
SPL2-mCherry
SPL3-mCherry
SPL4-mCherry
SPL5-mCherry
SPL6-mCherry
SPL8-mCherry
SPL7-mCherry
SRSF2-GFP
Analyte
Merge
SPL9-mCherry
SPL10-mCherry

**Fig. S4. Live cell confocal microscopy images of NCMC generated proteins targeted to nuclear speckles.** Live cell images of colon cancer (HCT-116) cells expressing SRSF2-GFP (Green) from the endogenous SRSF2 locus and induced expression of the indicated nucleolus targeted protein, scale 10 microns.

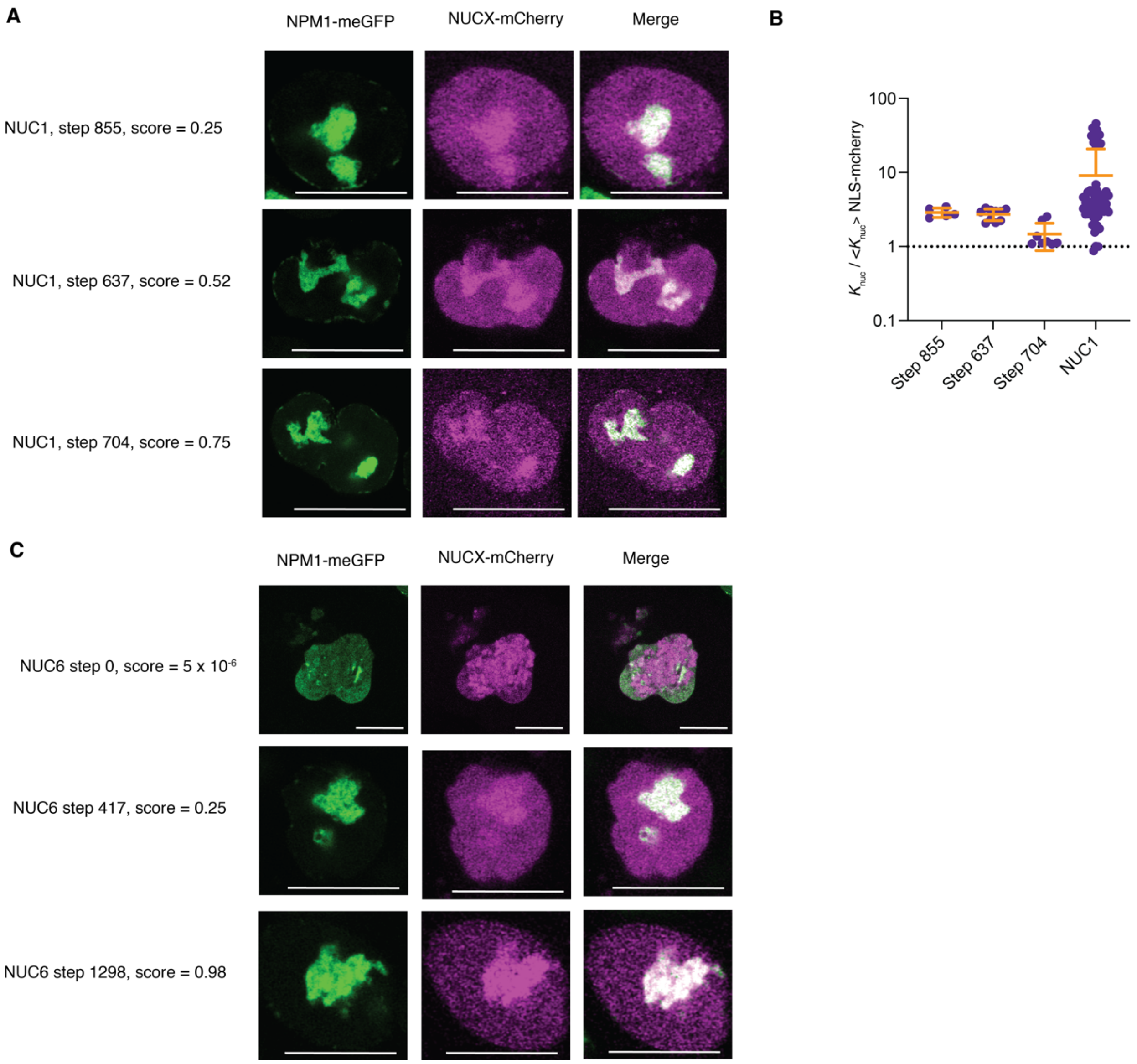


**Fig. S5.** Nucleolar partitioning and protein phenotype is sensitive to prediction strength. **A.** Live cell images of NUC12 proteins in colon cancer cells expressing NPM1-meGFP from the endogenous locus. Proteins were generated with MCMC and natural constraints across increasing scores. **B.** Quantification of the partition ratio of each NUC1-X step protein as compared to the average partition ratio of NLS-mCherry. **C**. Live cell images of NUC6 proteins in colon cancer cells expressing NPM1-meGFP from the endogenous locus, merged images are shown in panel Fig. 2E. Scale 10 microns. Quantification given in panel 2E.

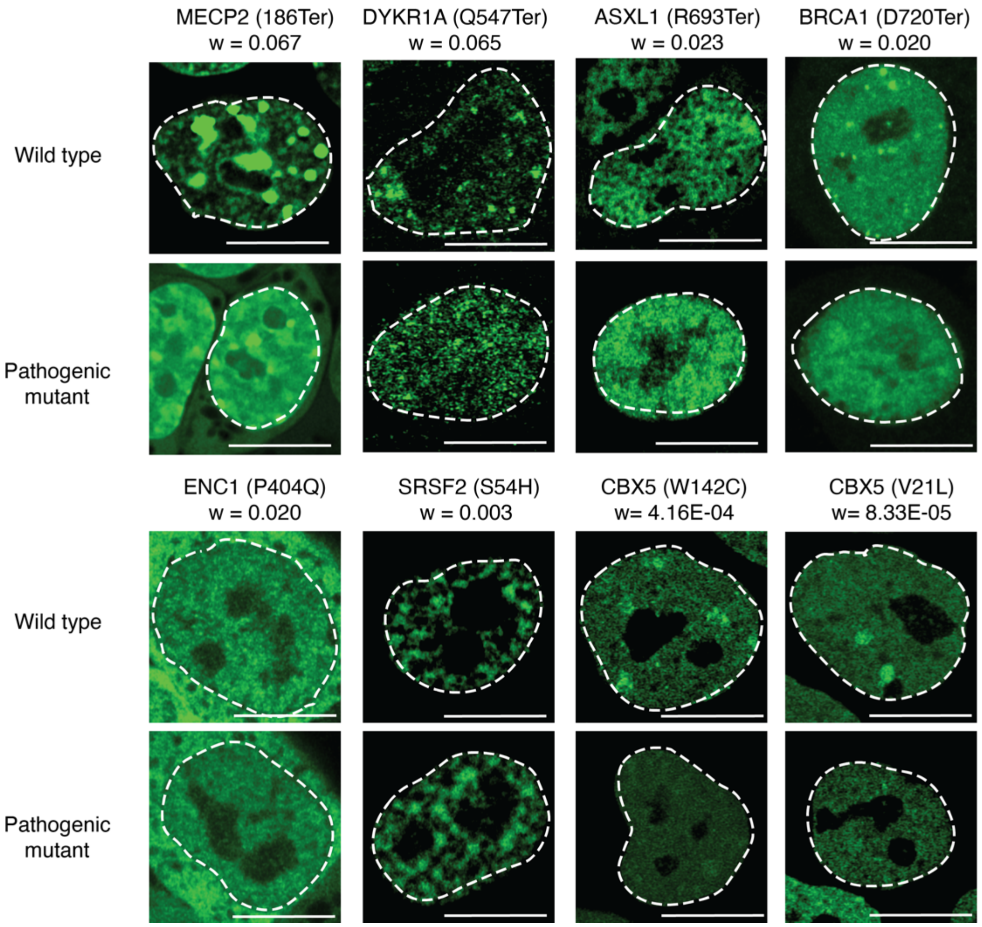


**Fig. S6. Live cell confocal micrographs of wild type and disease variants in mouse embryonic stem cells.** Live cell images of mouse embryonic stem cells (v6.5) expressing wild type and disease variant proteins from Table S3, scale 10 microns.
